# Immunoregulatory effects of Imuno TF^®^ (transfer factors) on Th1/Th2/Th17/Treg cytokines

**DOI:** 10.1101/2020.11.06.371435

**Authors:** Carlos Rocha Oliveira, Rodolfo Paula Vieira, Anderson de Oliveira Ferreira, Any Elisa de Souza Schmidt Gonçalves, Hudson Polonini

**Affiliations:** Anhembi Morumbi University, School of Medicine, Avenida Deputado Benedito Matarazzo, 6070, São Jose dos Campos-SP, Brazil, 12230-002; Federal University of Sao Paulo, Graduate Program in Sciences of Human Movement and Rehabilitation, Avenida Ana Costa 95, Santos-SP, Brazil, 11060-001; Universidade Brasil, Graduate Program in Bioengineering and in Biomedical Engineering, Rua Carolina Fonseca 235, Sao Paulo-SP, Brazil, 08230-030; Brazilian Institute of Teaching and Research in Pulmonary and Exercise Immunology (IBEPIPE), Rua Pedro Ernesto 240, Sao Jose dos Campos-SP, Brazil, 12245-520; Fagron BV, Lichtenauerlaan 182, 3062 ME, Rotterdam, The Netherlands; Infinity Pharma Brasil, Av. Pierre Simon de Laplace, 751 - Techno Park, Campinas - SP, 13069-320, Brazil

## Abstract

Transfer factors are known since 1955 due to their activities on the immune system. Although the reports on the effects on diverse immune mechanisms, their role on Th1, Th2, Th17 and Treg responses was still not described. In this sense, the present work focused on the evaluation of such immune responses. For that, human lymphocytes, and mice thymic, splenic and Peyer’s cells were stimulated with Lipopolysaccharides and Concanavalin A, and then treated with isolated transfer factors (Imuno TF^®^). The culture medium was harvested and the quantification of Th1 cytokines (IL-2 and IFN-γ), Th2 cytokines (IL-4, IL-5, and IL-13), Th17 cytokine (IL- 17), Treg cytokine (IL-35), inflammatory cytokines (IL-6 and TNF-α), and anti-inflammatory cytokine (IL-10) was performed, as well as the quantification of mRNA levels. Imuno TF^®^ positively regulated Th1 cytokines, while decreased Th2 cytokines. It also increased levels of mRNA and secretion of the anti-inflammatory cytokine IL-10, whereas it reduced levels of mRNA and the secretion of pro-inflammatory cytokines IL-6 and TNF-α. Finally, it reversed the hypersecretion of IL-17 and did not promote significant changes in IL-35 secretion. This highlights the role of Imuno TF^®^ in the regulation of the immune responses.

## 1. Introduction

Recently, it has been a wide interest in products that can strengthen the immune system against pathogenic microorganisms, such as bacteria and virus. One of these products is Imuno TF^®^, a nutritional supplement composed of oligo- and polypeptides fractions from porcine spleen, also known as transfer factors (TF). They were first described in 1955 and later characterized at the molecular level [1–3], being now understood as short chains of amino acids with small pieces of ribonucleic acid (RNA) attached [4,5].

To date, however, few studies have shed light on the fully elucidation of its mechanism of action. The current body of research shows that the TF main role in the immune system is the regulation of the immune responses. There is evidence that TF are released by CD4^+^ lymphocytes [6,7] and then essentially involved in the Th1 response, leading to an increase in Th1 cytokine levels (essentially IFN-γ) and suppressing the production of Th2 cells and their related cytokines such as IL-4, IL-5, IL-6, IL-10 and IL-13 [8,9].

Preliminary studies have also shown that TF can potentially play roles in: (i) CD4^+^ cells activation, triggering the production of IL-1 and IFN-γ [8–11]; (ii) macrophage activation, through the stimulation of IFN-γ production [9,10], (iii) modulation of the response to the recognition of micro-organism through the activation of TLR4-MD2 complex, which occurs through the MyD88-mediated NF-κB pathway [12]; (iv) inhibition of TNF-α production, through the inhibition of the NF-κB [12,13]; and (v) possible maintenance of physiological IL- 7 levels [14]. In this sense, TF can possibly regulate the immune system by stimulating it against microorganisms, while avoiding immune hyperresponsiveness.

In this regard, a current central issue is to understand which cytokines are indeed expressed and secreted when TF are endogenously released or exogenously administered. In this sense, the present work aims to help in the better understanding on the mechanisms of action of the TF on the immune system, focusing on Th1, Th2, Th17 and Treg responses.

## 2. Materials and Methods

### 2.1. Reagents

Dulbecco’s Modified Eagle Medium (DMEM), fetal bovine serum (FBS), penicillin-streptomycin, and phosphate-buffered saline (PBS) were obtained from Gibco BRL (Grand Island, CA, USA). 3-(4,5-Dimethylthiazol-2-yl)-2,5diphenyltetrazolium bromide (MTT), Lipopolysaccharides (LPS) and Concanavalin A (ConA) were purchased from Sigma-Aldrich (St. Louis, Mo., USA). Imuno TF^®^ was supplied by Infinity Pharma Brasil (Campinas, SP, Brazil), a Fagron company (Rotterdam, The Netherlands).

### 2.2. Animal and cell culture studies

The experiments were conducted using human lymphocytes and mice thymic, Peyer’s and splenic cells. This study was carried out after approval by the local Human and Animal Ethics Committees (approval numbers 4.142.254 and AN0018-2014, respectively). The animals were housed under specific pathogen-free conditions. The human lymphocytes were isolated from fresh heparinized venous blood via centrifugation over Ficoll gradients (Histopaque-1077; Sigma-Aldrich, Gillingham, United Kingdom) according to the manufacturer’s protocol. Human lymphocytes were incubated with LPS (1.0 *μ*g/mL) or ConA (2.5 μg/mL) for 2 hours, at 37° C, in a 5% CO2 humidified atmosphere and treated or not with Imuno TF^®^ (10 *μ*g/mL and 100 *μ*g/mL) for 24 hours. Mice were euthanized and their thymus, spleen, and Peyer’s patches from intestinal wall were removed for *in vitro* analysis. Purified cells obtained from mice tissues were washed and resuspended in appropriate culture medium. The cells were cultured in RPMI-1640, supplemented with 10% FCS, L-glutamine (2mol/L), and antibiotics (100U/mL penicillin and 100 mg/mL streptomycin). Five hundred thousand (5×10^5^) cells were seeded in 25 cm^2^ culture flasks and maintained at 37°C under an atmosphere of 5% CO_2_. Fresh medium was added every 2 days, and the cells were harvested and diluted 5-fold every 7 days.

Cells were stimulated with LPS (1.0 *μ*g/mL) or ConA (2.5 μg/mL) for 2 hours and treated or not with Imuno TF^®^ (10 *μ*g/mL and 100 *μ*g/mL) for 24 hours.

### 2.3. Thymocytes, Spleen and Peyer’s patches preparation

Male BALB/c mice, aged four-eight weeks, were obtained from the animal facility of the Federal University of São Paulo (UNIFESP) (São Paulo, Brazil). The mice thymic cells (thymocytes) were prepared by smashing thymic tissue through a 100 μm mesh. The cells were then washed and suspended in RPMI-1640 medium that was supplemented with 10% FCS, L-glutamine (2 mol/L), and antibiotics (100 U/mL penicillin and 100 mg/mL streptomycin). The cells were cultured in flasks at 37°C with 5% CO2 in humidified air. The spleen cells (2×10^6^ cells/well) from mice were incubated in culture flask and stimulated with LPS and Con A, as previously described. The cells were maintained in an incubator with 5 % CO2 at 37°C for 72 h. Subsequently, the cells were centrifuged at 700×g for 8 min and the supernatant was collected for cytokine determination by specific solid-phase sandwich enzyme-linked immunosorbent assay (ELISA). The BALB/C mice (4-8 weeks) had about 5 to 10 lymphoid follicles from the Peyer’s patches removed and transferred, under aseptic conditions, to a polyethylene tube containing DMEM medium supplemented with 10% foetal bovine serum, penicillin (100 U/mL) and streptomycin (100 μg/mL). The follicles were macerated, with the aid of a syringe plunger, against a nylon membrane with 40 μm pores. The cell suspension was centrifuged at 1000 rpm for 5 minutes. After centrifugation, the cells were washed twice with 5 ml of culture medium. At the end of the washing, the cell pellet was resuspended in 5 mL of supplemented medium, for counting viable lymphocytes, by staining with the trypan blue exclusion dye. The cells isolated from Peyer’s patches were transferred in triplicate, to a 24- well plate, at a concentration of 1.0×10^6^ cells/well. After treatment with Imuno TF^®^, stimulated or not with Con A or LPS, lymphocytes were incubated for 24 hours and the supernatant was used for cytokine determination by ELISA.

### 2.4. Cytotoxicity evaluation of Imuno TF^®^ by MTT assay

For all the experiments, a stock solution of Imuno TF^®^ was prepared by dissolving the compound directly in culture media at 10-300 μg/mL. The cell viability of control and Imuno TF^®^-treated human lymphocytes and mice thymocytes cells were measured using a standard MTT assay. Briefly, 5×10^4^ viable cells were seeded into clear 96-well flat-bottom plates in RPMI 1640 medium supplemented with 10% foetal bovine serum (FBS) and incubated with different concentrations (10-300μg/mL) of Imuno TF^®^ for 24h. Then, 10 μL/well of MTT (5 mg/mL) was added and the cells were incubated for 4h. Following incubation, 100μL of 10% sodium dodecyl sulphate (SDS) solution in deionized water was added to each well and left overnight. The absorbance was measured at 595 nm using a microplate reader (Molecular Device).

### 2.5. Cytokines quantification

Cells were stimulated for 2 hours with LPS or Con A, treated or not with Imuno TF^®^ (100μg/mL) for 24h, as describe above (item 2.2). The culture medium was harvested and the quantification of Th1 cytokines (IL-2 and IFN-γ), Th2 cytokines (IL-4, IL-5, and IL-13), Th17 cytokine (IL-17), Treg cytokine (IL-35), inflammatory cytokines (IL-6 and TNF-α), and antiinflammatory cytokine (IL-10) was performed using ELISA kits (R&D Systems, Minneapolis, MN, USA) according to the manufacturer’s protocol.

### 2.6. Reverse transcription quantitative PCR (RT qPCR)

Total RNA extracted from cells samples was converted to cDNA using a SuperScript^®^ III RT kit (Invitrogen, Carlsbad, CA), according to the manufacturer’s protocol. The concentration of RNA was detected using a NanoDrop 2000 (Thermo Fisher Scientific, Inc.). GAPDH were used as the internal control. The thermocycling conditions were as follows: 95°C for 10 min followed by 35 cycles of 95°C for 15 sec and 55°C for 40 sec. The 2 ΔΔCq method was used to quantify the relative gene expression levels of the target genes. Relative standard curves were generated by serial dilutions and all samples were run in triplicates. **Table 1** indicates the sense and antisense sequences of primers used in qRT-PCR analysis.

**Table 1:**
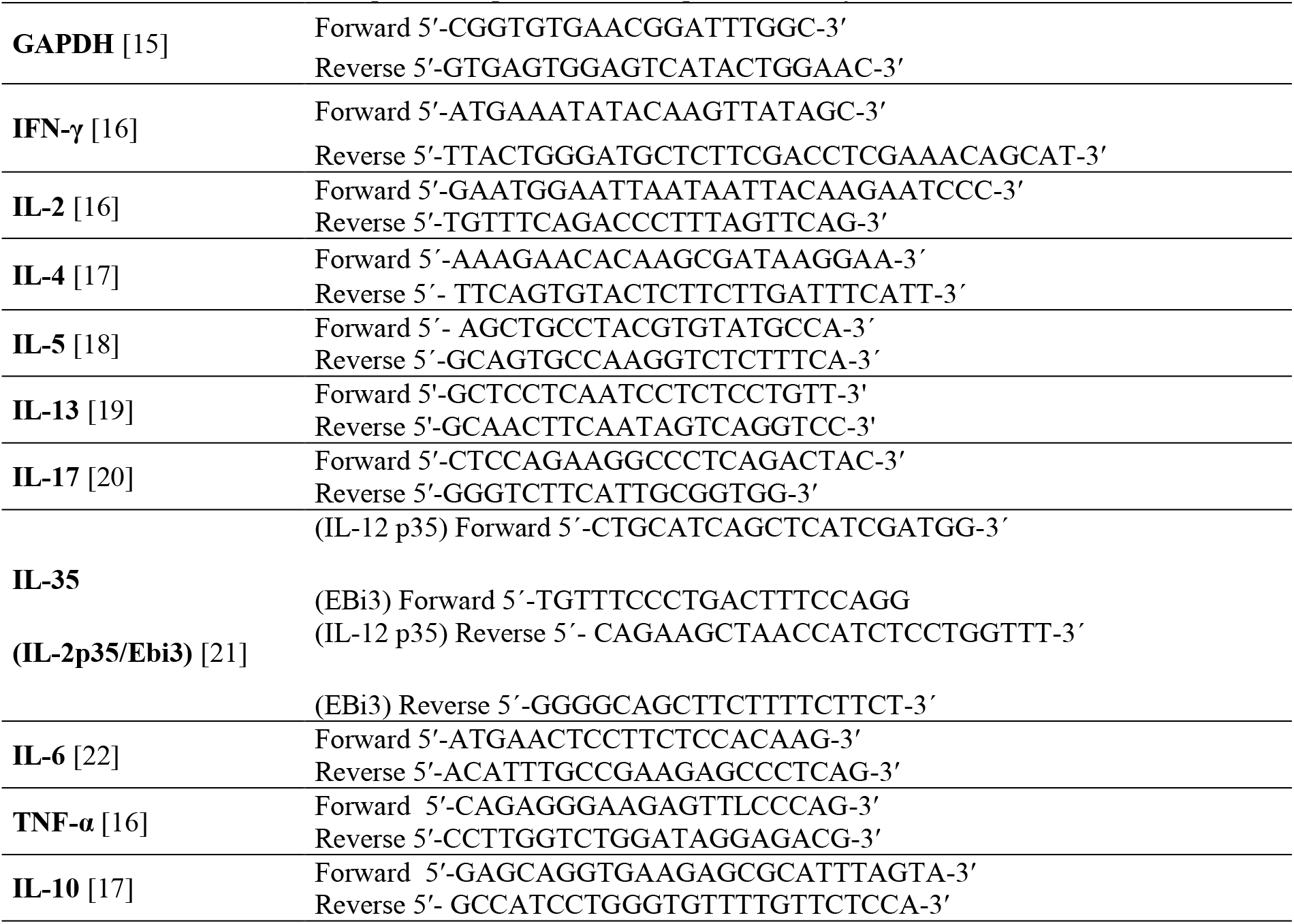
Sense and antisense sequences of primers used in qRT-PCR analysis.

### 2.7. Statistical analysis

The results were expressed as the mean ± standard error of mean (SEM) from at least three independent experiments, unless stated otherwise. Paired data was evaluated by Student’s t-test. One-way analysis of variance (ANOVA) was used for multiple comparisons. A *p* value of <0.05 was considered significant.

## 3. Results

### 3.1. Cell viability

To determine the optimal concentration without compromising cell viability, the MTT test with dose-response of Imuno TF^®^-mediated effects on cell proliferation was performed (**Figure 1**). The concentrations of 10 μg/mL and 100 μg/mL were defined as the study doses.

**Figure 1:**
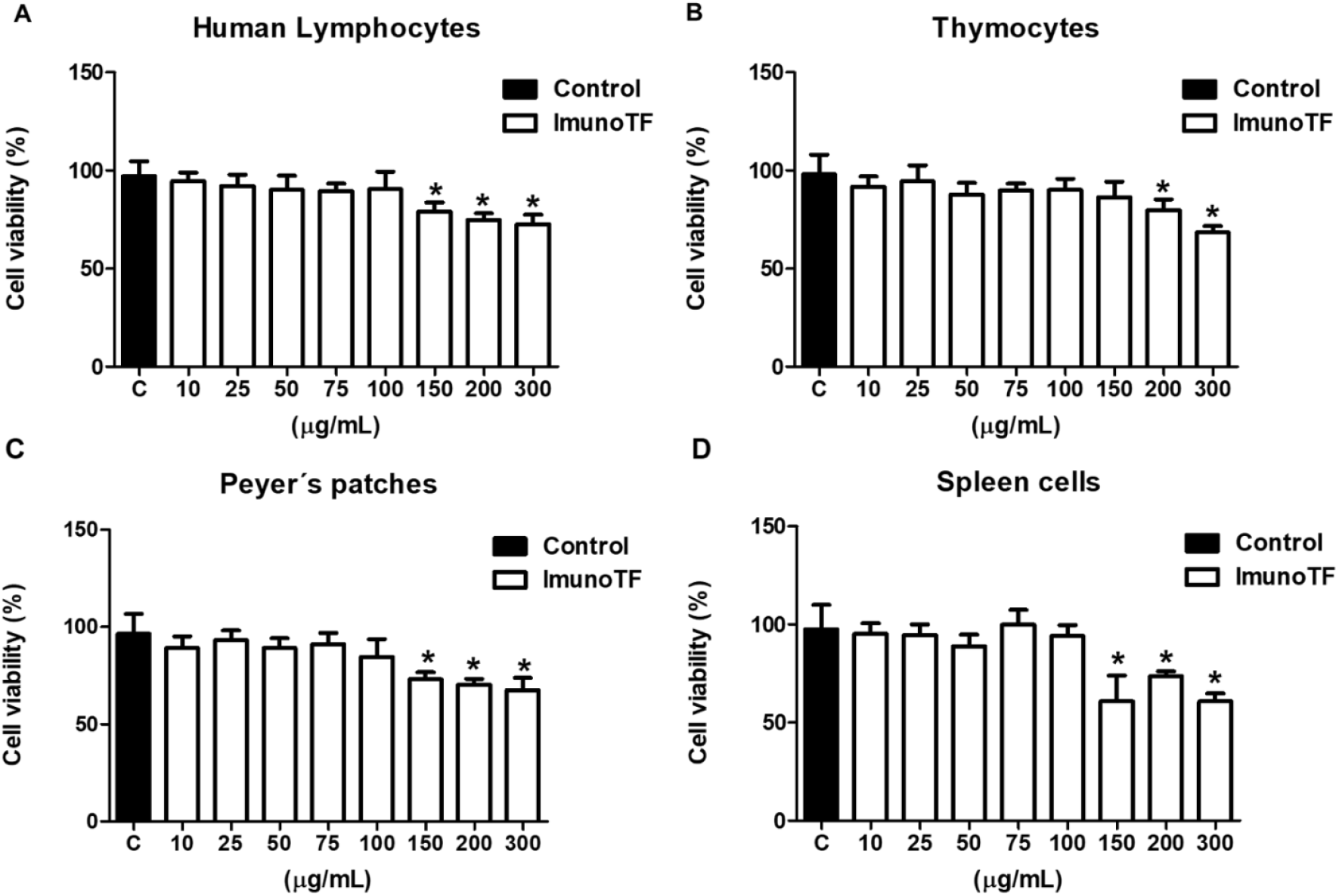
Effects of Imuno TF^®^ on (A) human lymphocytes, (B) murine thymocytes, (C) murine Peyer’s cells and (D) murine splenocytes. Cells were treated with different concentrations of Imuno TF^^®^^ for 24 hours; 10 and 100 μg/mL were defined as test concentrations for all experiments. Values are expressed as mean ± SEM (n=3), and * indicates *p*< 0.05.

### 3.2. Th1 cytokines secretion

**Figure 2** shows the significant increase of IL-2 and IFN-γ levels (*p<0.05) in the supernatant of human lymphocytes and murine thymocytes (Imuno TF^®^ standalone or in the presence of LPS and ConA). In the lymphocytes, treatment with Imuno TF^®^ demonstrated a reduced cytokines secretion when compared to LPS and ConA groups, indicating that Imuno TF^®^ does not compromise the Th1 response. Thymocytes also showed reduced levels of IL-2 and IFN- γ, but with values similar to control group and statistically lower (# p <0.05) than LPS and ConA-treated cells.

**Figure 2:**
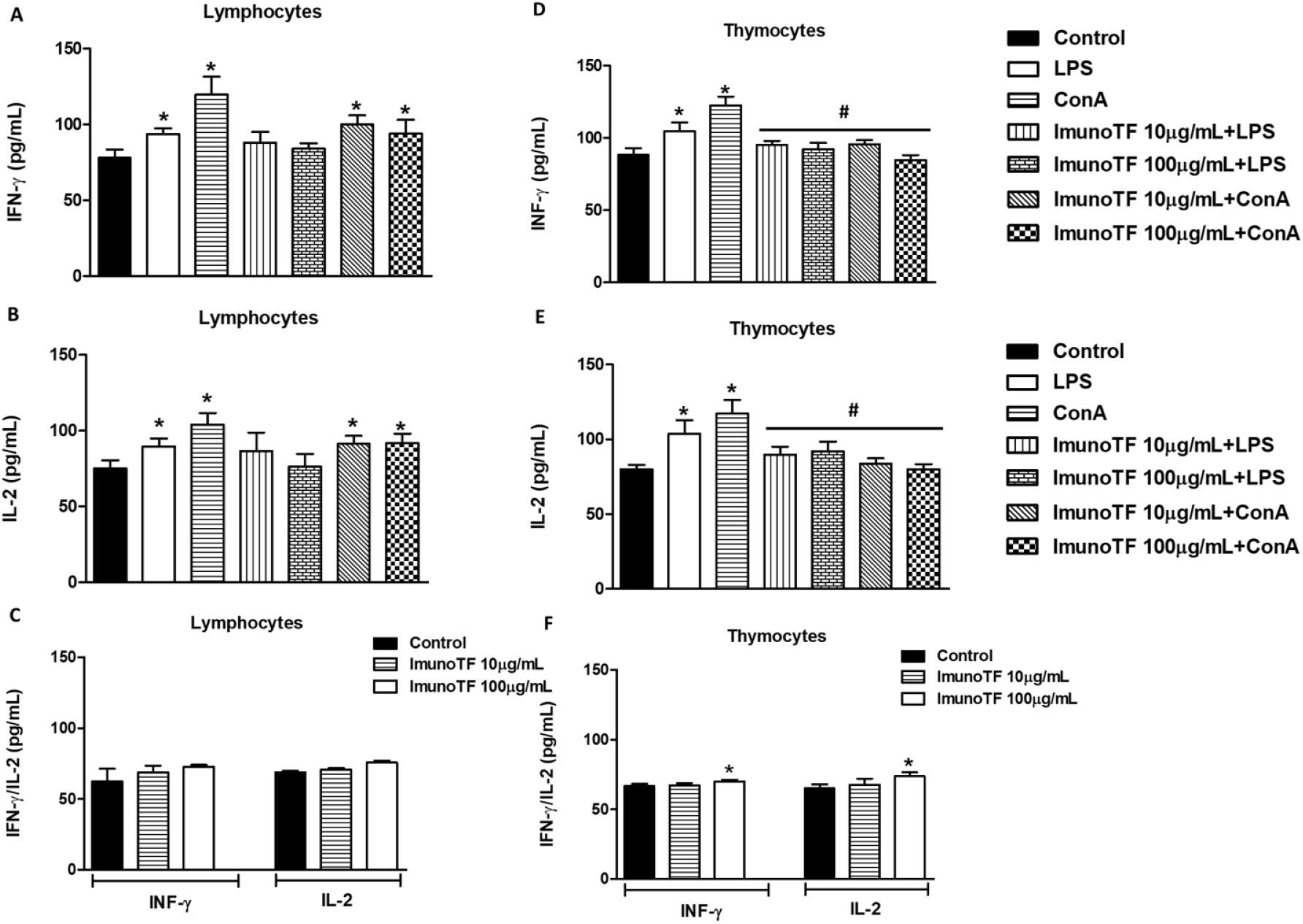
Evaluation of the secretion of Th1 cytokines by human lymphocytes (A and B) and murine thymocytes (D and E). The concentrations of 10 and 100 μg/mL of Imuno TF^®^ were used for 24 hours. Values are expressed as mean ± SEM (n=3), * indicates p < 0.05 vs. control (non-treated cells), and # indicates p < 0.05 vs. LPS and ConA-treated cells. (C and D) show the influence of Imuno TF^®^, alone, on the secretion of Th1 cytokines.

In addition, **Figure 3** shows the secretion of IL-2 and IFN-γ on Peyer’s patches cells and on splenocytes, both from mice. The results showed a similar secretion profile when compared with human lymphocytes.

**Figure 3.**
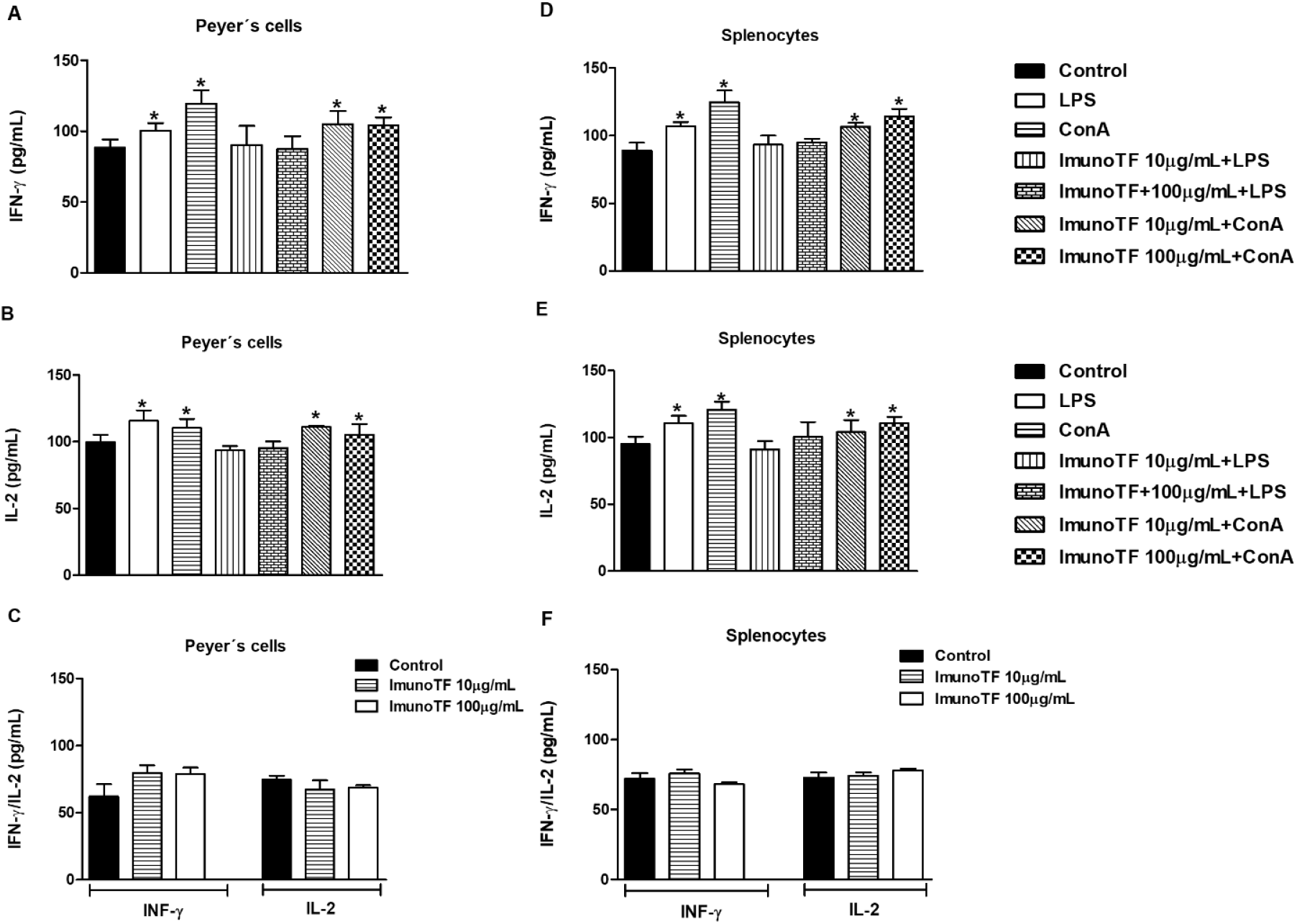
Effects of Imuno TF^®^ on IL-2 and IFN-γ secretion by Peyer’s cells (A and B) and splenocytes (D and E). Cells were treated with 10 and 100 μg/mL of Imuno TF^®^ for 24 hours. Values are expressed as mean ± SEM (n=3), * indicates p < 0.05 vs. control (non-treated cells). (C and F) show the influence of Imuno TF^®^ on the secretion of Th1 cytokines.

### 3.3. Th2 cytokines secretion

Differently from the profile observed in Th1 cytokine secretion, Th2 cytokines were reduced on the cells that were challenged by LPS and ConA and were treated with Imuno TF^®^ (#p <0.05). In addition, Imuno TF^®^, in the absence of LPS and ConA, significantly reduced (*p <0.05) the cytokine secretion of the Th2 response. Figure 4 shows the influence of Imuno TF^®^ on cytokine secretion.

**Figure 4:**
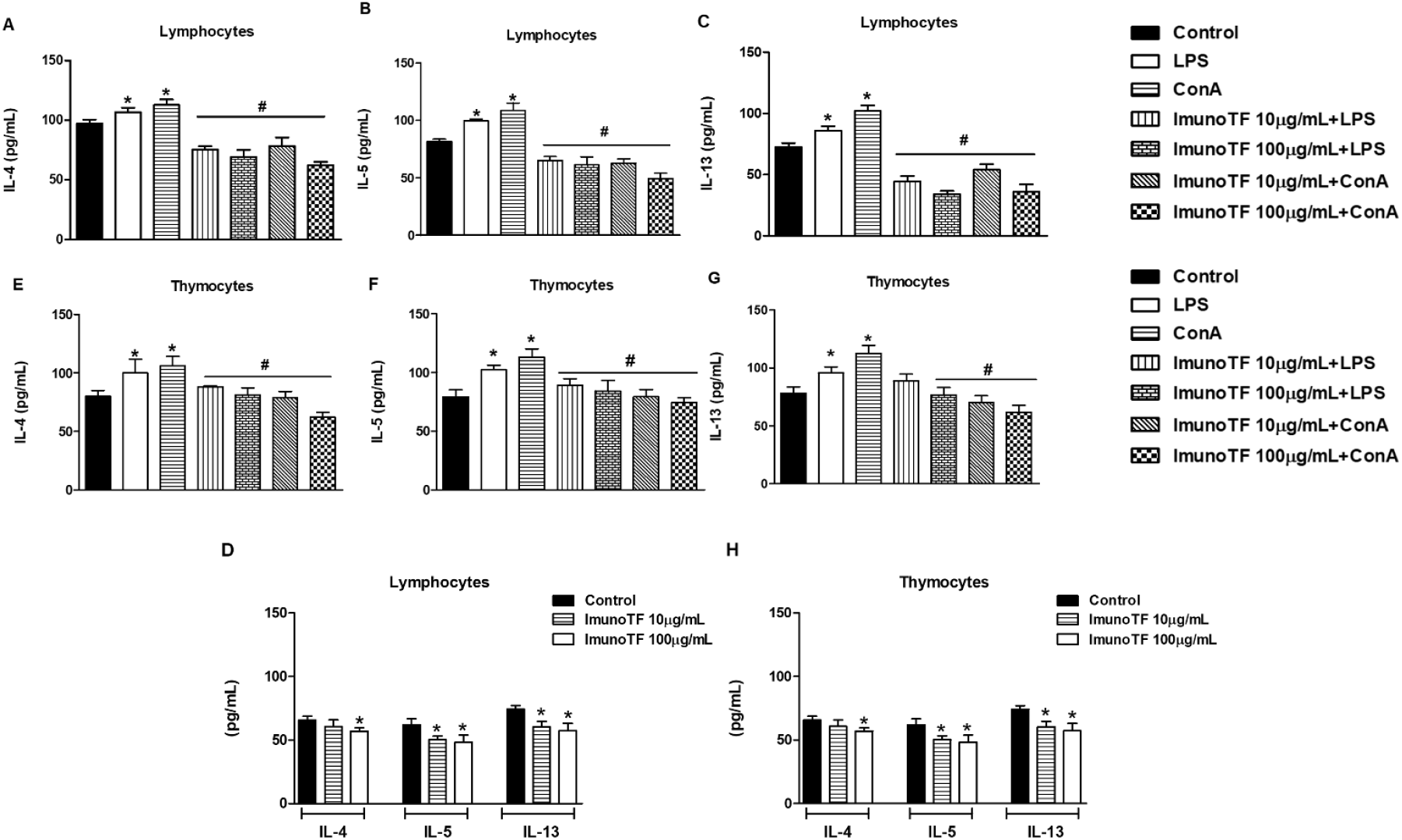
Th2 cytokines secretion by human lymphocytes and murine thymocytes. Cells were treated with 10 and 100 μg/mL of Imuno TF^®^ for 24 hours. Values are expressed as mean ± SEM (n=3), * indicates p < 0.05 vs. control (non-treated cells) and # indicates p < 0.05 vs. LPS and ConA-treated cells. (D and H) show the influence of Imuno TF^®^ on the secretion of Th2 cytokines.

The influence of Imuno TF^®^ alone on the cytokine secretion of Peyer’s patches and the splenocytes obtained both from mice was also evaluated (**Figure 5**).

**Figure 5:**
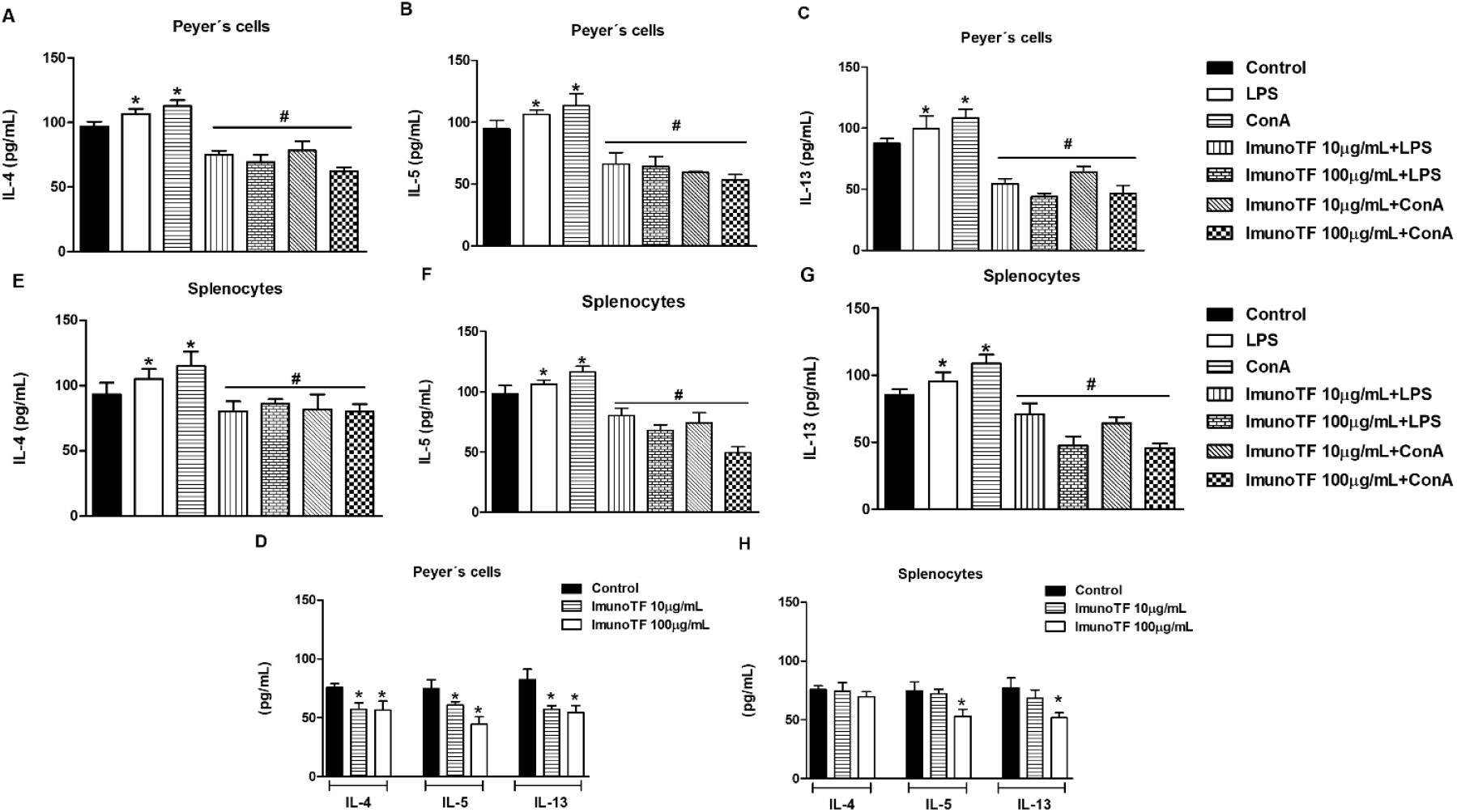
Secretion of IL-4, IL-5, and IL-13 secretion by Peyer’s cells (A, B, and C) and splenocytes (E, F, and G) after treatment with Imuno TF^®^ on. Cells were treated with 10 and 100 μg/mL for 24 hours. Values are expressed as mean ± SEM (n=3), * indicates p<0.05 vs. control (non-treated cells) and # indicates p<0.05 vs. LPS and ConA-treated cells. (D and H) show the influence of Imuno TF^®^ on the secretion of Th2 cytokines.

### 3.4. Th17 and Treg secretion

The secretion of IL-17 and IL-35 were evaluated and are represented on **Figures 6–7**. Regarding on human lymphocytes, the treatment of cell cultures with Imuno TF^®^ reduced the levels of IL-17 (#p <0.05), when compared with stimulated-cells (LPS and ConA). The secretion of IL-35 showed no significant reduction. On the other hand, the Imuno TF^®^-treated thymocyte cultures showed significant reduction (#p <0.05) in IL-35 secretion (**Figure 6**).

**Figure 6:**
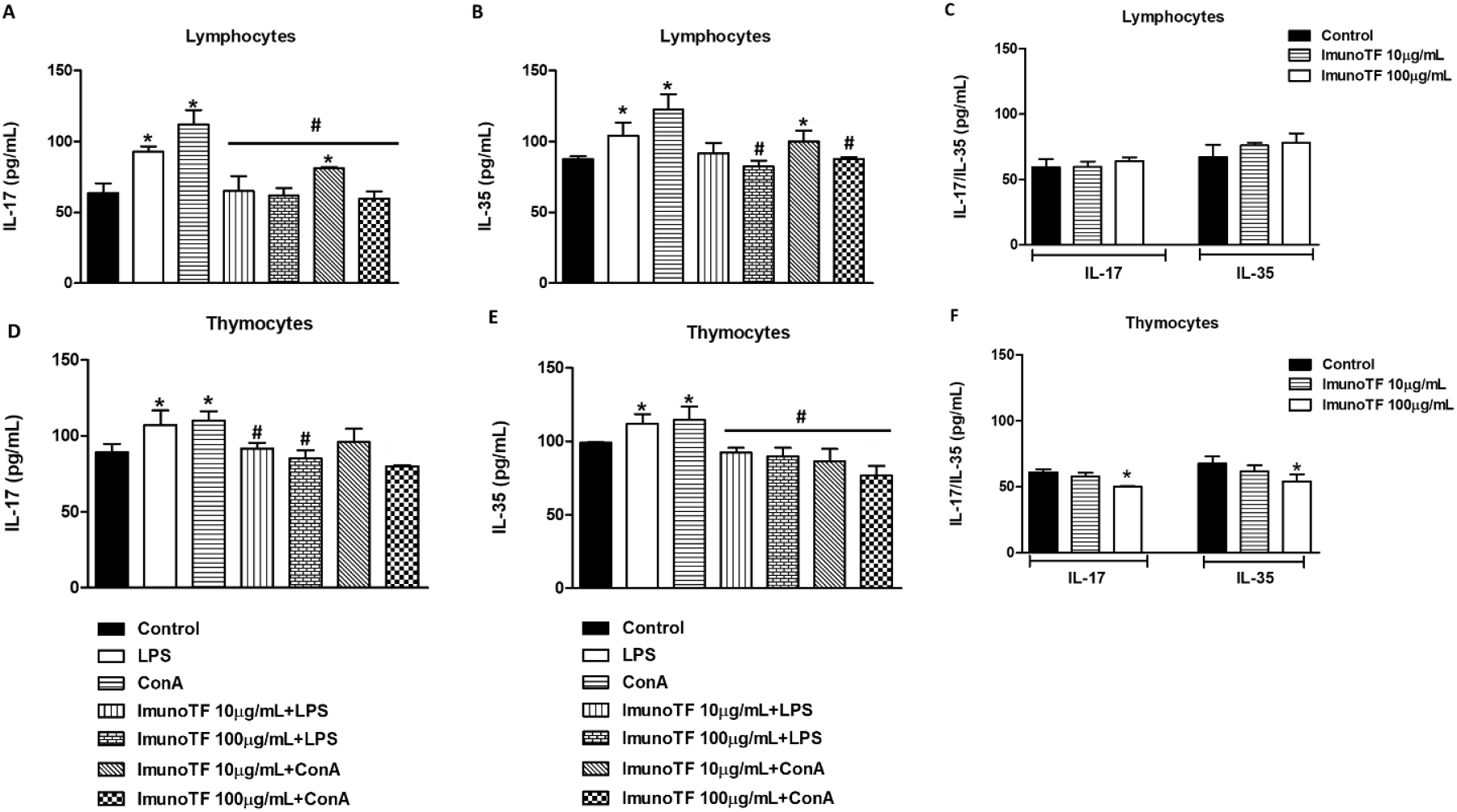
Effects of Imuno TF^®^ on IL-17 and IL-35 from human lymphocytes (A and B) and murine thymocytes (D and E). Cells were treated with 10 and 100 μg/mL of Imuno TF^®^ for 24 hours. Data shown are representative of three independent experiments. The values are expressed as mean ± SEM, *p < 0.05 indicates statistical difference vs. control (non-treated cells), and #p < 0.05 indicates statistical difference vs. LPS and ConA-treated cells. Figures C and F show the influence of Imuno TF^®^ on the secretion of IL-17 and IL-35.

The secretion of IL-17 and IL-35 in Peyer’s cells and splenocytes, showed a significant reduction only in the secretion of IL-17 dosed in Peyer’s cells treated with Imuno TF^®^ (#p <0.05). Specifically, Peyer’s cells and splenocytes stimulated with LPS showed a reduction in IL-35 (**Figure 7B**) and in IL-17 (**Figure 7D**).

**Figure 7.**
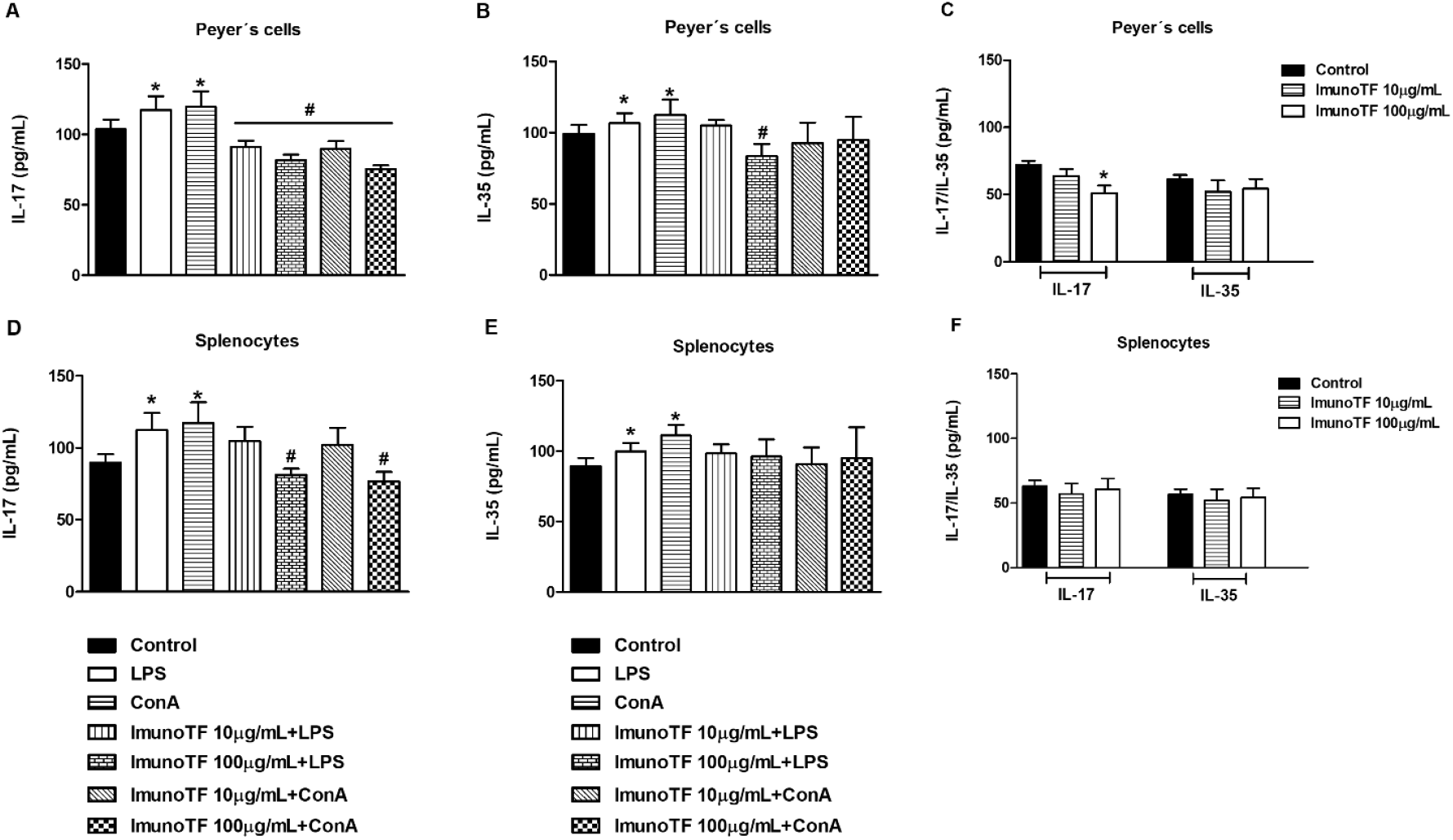
Effects of Imuno TF^®^ on IL-17 and IL-35 from murine cells. Peyer’s cells (A and B) and splenocytes (D and E). Cells were treated with 10 and 100 μg/mL of Imuno TF^®^ for 24 hours. Data shown are representative of three independent experiments. The values are expressed as mean ± SEM, *p < 0.05 indicates statistical difference vs. control (non-treated cells), and #p <0.05 indicates statistical difference vs. LPS and ConA-treated cells. Figures C and F show the influence of Imuno TF^®^ on the secretion of IL-17 and IL-35.

### 3.5. Inflammatory and anti-inflammatory cytokines

We also investigated the role of Imuno TF^®^ in the secretion of inflammatory cytokines (IL-6 and TNF-α) and IL-10, a cytokine - with multiple effects in immunoregulation. The results in **Figure 8**, showed that TNF-α values were reduced (#p <0.05) in the presence of Imuno TF^®^, when compared to the secretion values of cells stimulated with LPS and ConA. Regarding IL- 6, only in the treatment situations with the highest concentration of Imuno TF^®^ (100μg/mL), we observed a significant reduction in the secretion of IL-6, when compared with stimulated cells.

The secretion of IL-10, a cytokine with anti-inflammatory properties, is indicated on **Figure 9**. In all situations of stimulus with LPS and ConA, no significant changes were observed. Except in human lymphocytes, where IL-10 secretion was significantly lower (#p <0.05), when compared to cells stimulated with LPS and ConA (**Figure 9A**). The results of the evaluation of the Imuno TF^®^ alone (without stimulation of LPS or ConA), showed a significant increase in the secretion of IL-10, with the highest concentration tested (**Figure 9E–9G**). However, **Figure 9H** shows that in splenocytes, no significant changes were observed in any treatment situation for these cells.

**Figure 8:**
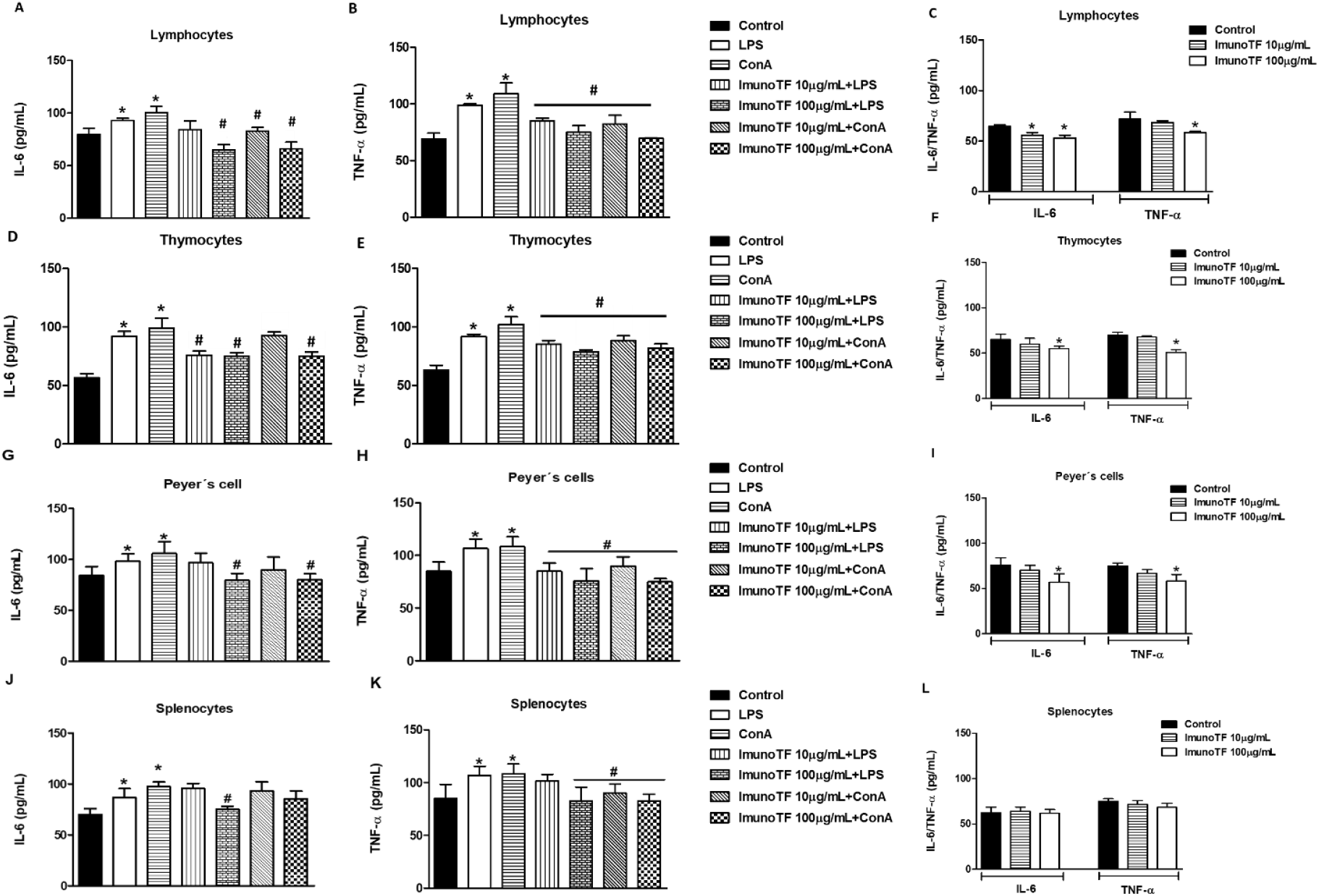
Effects of Imuno TF^®^ on IL-6 and TNF-α from human lymphocytes and murine cells (thymocytes, splenocytes and Peyer’s cells). Lymphocytes cells (A, B and C); Thymocytes (D, E and F); Peyer’s cells (G, H and I) and Splenocytes (J, K and L). Cells were treated with 10 and 100 μg/mL of Imuno TF^®^ for 24 hours. Data shown are representative of three independent experiments. The values are expressed as mean ± SEM, *p < 0.05 indicates statistical difference vs. control (non-treated cells), and #p < 0.05 indicates statistical difference vs. LPS and ConA-treated cells. (C, F, I and L) show the influence of Imuno TF^®^ on the secretion of on IL-6 and TNF-α.

**Figure 9:**
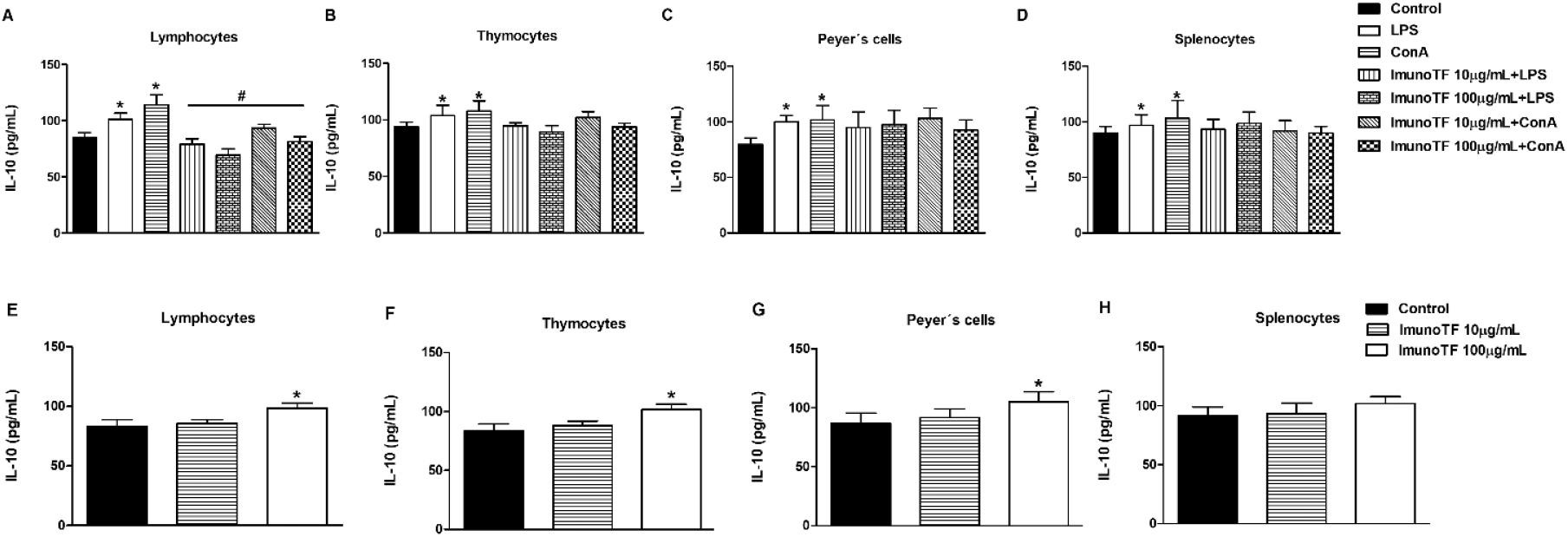
Effects of Imuno TF^®^ on IL-10 from human lymphocytes (A) and murine cells (B, C and D). Cells were treated with 10 and 100 μg/mL of Imuno TF^®^ for 24 hours. Data shown are representative of three independent experiments. The values are expressed as mean ± SEM, *p < 0.05 indicates statistical difference vs. control (nontreated cells), and #p < 0.05 indicates statistical difference vs. LPS and ConA-treated cells. (E-H) show the influence of Imuno TF^®^ on the secretion of on IL-10.

### 3.6. Quantification of mRNA levels by RT-PCR of Th1, Th2, Th17, Treg, Inflammatory and anti-inflammatory cytokines

The influence of Imuno TF^®^ on the modulation of mRNA levels of the evaluated cytokines is described in **Figure 10**. Except for IFN-γ, in Peyer’s cells, in all other evaluations we can observe a significant increase (* p <0.05) or an increase trend, in relation to the cytokines of the Th1 response (**Figure 10A, 10E, 10I, and 10N**). The cytokines of the Th2 response are essentially altered in human lymphocytes, when most of them were significantly reduced. In other types of cells, one point or the other showed a significant reduction, although the tendency of reduction is observed. The **Figure 10** (**10D**, **10H**, **10M**, and **10Q**) indicates that all cells evaluated (murine or human lymphocytes), showed a significant increase in IL-10 (*p <0.05). Finally, on inflammatory cytokines, IL-6 was significantly reduced (*p<0.05) in all situations where the highest dose of Imuno TF^®^ (100μg/mL) was used, except for splenocytes (**Figure 10P**).

**Figure 10.**
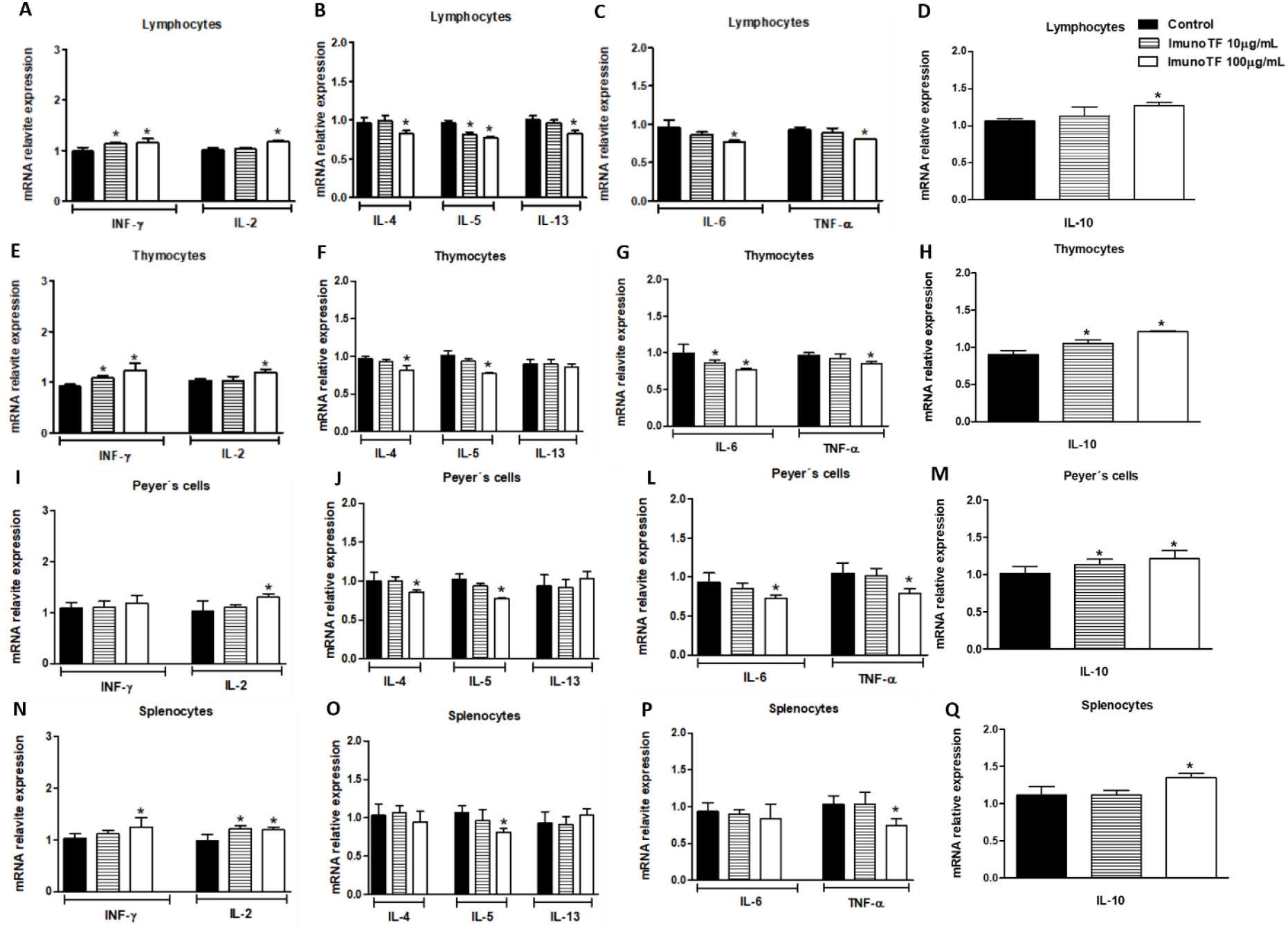
Levels of mRNA by RT-PCR of Th1 and Th2 response, IL-17, IL-35, inflammatory and anti-inflammatory cytokines. Effects of Imuno TF^®^ (10 and 100 μg/mL) on mRNA levels of cytokines from human lymphocytes and murine cells (thymocytes, Peyer’s cells and splenocytes). Cells were treated for 24 hours. Data shown are representative of three independent experiments. The values are expressed as mean ± SEM, and *p < 0.05 indicates statistical difference vs. control (non-treated cells).

## 4. Discussion

TF have been described in the last decades as natural, small peptides of <10 kDa in molecular weight. Reports from literature show their diverse applications in clinical conditions where the immune system plays a role, such as varicella-zoster in children with acute leukaemia, HIV, hepatitis C, and chronic mucocutaneous candidiasis [23–27].

The present study focused on Imuno TF^®^, an ultrafiltered extract from porcine spleen containing TF, in the modulation of inflammatory markers associated with Th1, Th2, Th17 and Treg responses, as well as in relation to pro- and anti-inflammatory cytokines, such as IL- 6, and TNF-α and IL-10, respectively. As previously described, human lymphocytes and mice cells (thymocytes, Peyer’s cells and splenocytes) were used, and the cell viability assay showed that only concentrations greater than 100 μg/mL of Imuno TF^®^ reduced the viability of the cell types of this study, showing alignment with safety data described previously [9,28]. Additionally, two mitogens were used to evaluate the immunomodulatory effects of the Imuno TF^®^: LPS and Con A. Both LPS and Con A are widely used in in vitro and in vivo tests to assess the ability of cells to secrete various types of cytokines [29–31].

We observed that the Imuno TF^®^ (at 10 μg/mL and 100 μg/mL) reduced the secretion of cytokines related to the Th1 response, which were significantly increased in the presence of LPS and ConA. However, when comparing Th1 cytokine secretion (IL-2 and IFN-γ) with nontreated cells, the reduction is not significant. In addition, we observed that Imuno TF^®^, in the absence of LPS and Con A, did not negatively modulate Th1 cytokine secretion, suggesting that it could, at least in part, positively influence the determination of a Th1 response. These results corroborate previous reports from the literature that suggested that the TF’s mechanism of action would be the increase of Th1 response and consequent decrease of Th2 cytokines such IL-4, IL-5, and IL-13 [8,9,32].

Thus, to verify the signalling of the Th2 response, we assessed the secretion of their related cytokines (IL-4, IL-5, and IL-13). The results showed that the treatment with Imuno TF^®^ reduced the secretion of these cytokines both in LPS- and ConA-stimulated cells and non-stimulated cells, which also is aligned with the hypothesis of Th1-response stimulation and Th2-response reduction. In addition to the secretion of Th1 and Th2 cytokines, the mRNA levels of these cytokines were evaluated. The results obtained were compatible with the secretion of each cytokine, showing that the mechanism of genetic transcription may have been activated and/or repressed by Imuno TF^®^.

The effects of Imuno TF^®^ on secretion and mRNA levels of pro and anti-inflammatory cytokines were also evaluated in this study. Pro-inflammatory cytokines (i.e. IL-6 and TNF-α) are involved in the up-regulation of inflammatory reactions [33]. Our results showed that both the secretion and the mRNA levels of the pro and anti-inflammatory cytokines were altered in the presence of Imuno TF^®^: the secretion of the pro-inflammatory cytokines IL-6 and TNF-α in cells stimulated with LPS and Con A were reduced. Isolated, Imuno TF^®^ also significantly reduced the secretion of IL-6 and TNF-α and the levels of their respective mRNA. Murine splenocytes were the only cells that did not show a significant reduction in IL-6, both in secretion and in mRNA levels. These results observed in splenocytes may be related to the fact that we used spleens from young animals – for instance, Park and collaborators [34] showed that the production of IL-6 in the mouse spleen was elevated with aging and that IL-6 production by stromal cells played a major role in this enhancement. Thus, the potential of Imuno TF^®^ in reducing pro-inflammatory cytokines suggests that this TF plays a role avoiding immune hyperresponsiveness and hyperinflammatory condition, which corroborates reports from literature [12,35].

In contrast to the negative modulation of pro-inflammatory cytokines observed with the treatment of human and murine cells with Imuno TF^®^, both the secretion and the levels of IL- 10 mRNA did not present values significantly reduced after stimulation with LPS and Con A, with the exception of human lymphocytes, whose values were significantly reduced. Without stimulation of the mitogens used, Imuno TF^®^ positively modulated the mRNA levels and the secretion of IL-10. These results corroborate the proposed mechanism of action in which releasing IL-10, an inhibitory cytokine from Th2 cells that plays a vital inhibiting induction of pro-inflammatory cytokines that could lead to the development of autoimmune disorders [9]. Additionally, evidences suggested that the production of IL-10 is associated with Treg cells [36,37].

Finally, Th17 and Treg responses were also investigated. Th17 cells stimulate cells to recruit neutrophils to sites of infection, mediating immune responses against microorganisms, while Treg cells inhibit immune responses to maintain immune homeostasis [38]. In cells stimulated with LPS and Con A, IL-17 was reduced with the treatment of Imuno TF^®^. In non-stimulated cells, IL-17 secretion did not change, except for thymocytes and Peyer’s cells, where the highest concentration of Imuno TF^®^ (100 μg/mL) significantly reduced the secretion of IL-17. The Th17 cells are responsive to pathogens as well as to the commensal gut microbiota. While they have been associated with immune protection against pathogens they are also known to drive autoimmune inflammation in the gut [39]. As IL-17 is a pro-inflammatory cytokine that plays critical roles in host defence against extracellular bacteria and fungi and also in the pathogenesis of autoimmune diseases [40]. In this sense, the reduction in the secretion of IL- 17 by Peyer’s cells could indicate a discreet, benefit of Imuno TF^®^ in the intestinal microbiota’s homeostasis. In addition, regarding thymocytes, recent studies in human autoimmune diseases and in animal models have indicated that IL-17 may be the essential T cell cytokine in the autoimmune process mediated by these cells [41], suggesting that Imuno TF^®^ could assist in the treatment or prophylaxis of autoimmune diseases. On the other hand, our results did not indicate significant changes in the secretion of IL-35 in human or murine cells, except for the population of thymocytes (significant reduction). Although IL-35 is not constitutively expressed in tissues, it can suppress inflammatory responses of immune cell by its selective activity on different T-cell subsets [42,43]. The absence of influence of Imuno TF^®^ on the secretion of IL-35 allows us to ratify, at least partially, the Th1 activation of the product.

## 5. Conclusions

Our results showed that the Imuno TF^®^ regulated both the secretion and mRNA levels of cytokines of the Th1 and Th2 responses. Imuno TF^®^ negatively regulated the cytokines of the Th2 response, and positively regulated the cytokines of the Th1 response. In addition, Imuno TF^®^ increased levels of mRNA and secretion of the anti-inflammatory cytokine IL-10, whereas it reduced levels of mRNA and the secretion of pro-inflammatory cytokines IL-6 and TNF-α. Finally, the results showed that Imuno TF^®^ reversed the hypersecretion of IL-17 triggered by LPS and ConA stimuli and did not promote significant changes in IL-35 secretion.

## Data Availability

All data are available upon request.

